# Metabolomic-Genomic prediction drastically improves prediction accuracy of breeding values in crop breeding

**DOI:** 10.1101/2022.08.03.502591

**Authors:** Xiangyu Guo, Pernille Sarup, Ahmed Jahoor, Just Jensen, Ole Fredslund Christensen

## Abstract

Metabolomics is intermediate stage between genotype and phenotype, and therefore useful for breeding. Objective was to investigate heritabilities and accuracies of genetic evaluation of malting quality (MQ) traits by integrating both genomic and metabolomic information. In total, 2,430 plots of 562 malting spring barley lines from three years and two locations were included. Five MQ traits were measured in wort produced from each plot. Metabolomic features used were 24,018 Nuclear Magnetic Resonance intensities measured on each wort sample. Methods for statistical analyses were genomic best linear unbiased prediction (GBLUP) and metabolomic-genomic best linear unbiased prediction (MGBLUP). Accuracies of predicted breeding values were compared using two cross-validation strategies: leave-one-year-out (LOYO) and leave-one-line-out (LOLO). Plot heritabilities ranged from 0.060 to 0.279 for GBLUP, and increased to range from 0.123 to 0.283 for MGBLUP. LOLO scheme yielded higher accuracies than LOYO scheme, and MGBLUP yielded higher accuracies than GBLUP regardless of cross-validation scheme.

**Author Summary:** Metabolomics is intermediate stage between genotype and phenotype, and therefore useful for breeding. We carried out a study on 2,430 plots of 562 malting spring barley lines and 24,018 Nuclear Magnetic Resonance intensities measured on each wort sample, to investigate the prediction of breeding values of five malting quality traits by integrating genomic and metabolomic information using a metabolomics-genomic model. Results showed an increase in prediction accuracy for this model compared to a genomic model.

## Introduction

Plant breeding activities aim to maximize selection gain [1]. To reach this goal, scientific breeding activities aim at early and accurately ranking the selection candidates by their genetic merit, estimated through proper experimental design and analysis of their performance [2]. The history of scientific breeding of plants can be traced back to the theoretical basis of Mendel’s inheritance laws and the concepts of evolutionary biology [1]. The rediscovery of Mendelian inheritance formed the basis for quantitative breeding of plants [3], and following that, many quantitative genetic techniques have become common practice in plant breeding [4]. In practical breeding, measurements of phenotypes are used to inform selection, and generally these measurements must be conducted in well-designed experiments repeated over locations and years to avoid confounding effects from the environment when selection is based on the phenotype alone [2, 5]. However, this traditional phenotypic selection is time-consuming and expensive, which limits the efficacy of the breeding program. Genomic selection (GS) has shown to be a promising strategy for increasing genetic gain in plant breeding [6]. In the procedure of GS, dense genome-wide markers and phenotypes from a training population are used to predict genomic breeding values of candidates in a test population with only genotype but not phenotype [7]. A prerequisite for practical use of GS is the availability of genome-wide markers on many individuals at a low cost [8]. The application of GS makes early selection before the collection of phenotypes possible, which can increase the efficiency of the breeding efforts considerably due to shorter generation intervals and larger populations available for selection.

Similar to dense genome-wide DNA-markers, omics data such as transcriptomic and metabolomic data are becoming available in increasingly larger quantities and at decreasing cost. Such omics data consist of measurements of effects that are intermediate between genotype and phenotype of interest breeding. In recent years, the integration of multi-omics data has been of great interests to predict traits phenotypes as reported by several studies [9–11], in which multi-omics phenotypic prediction has been shown as promising methods. Prediction of phenotypes is a prediction of both genetic and environmental effects. However, only predicted genetic effects are of interest for breeding purposes. Accurate prediction of breeding values is crucial in plant breeding programs to maximize genetic gain. In order to meet the need for integrating various multi-omics data resources in a genetic evaluation system, a joint model for phenotypes and omics data was developed by Christensen et al. [12], where the computation of predicted breeding value from such a model was also described. The methods provide great opportunity for prediction of breeding values in practical plant or animal breeding, especially when the traits of interests are costly to measure or difficult to improve.

Barley is the most common source for malt used in brewing alcoholic beverages [13]. Malting quality (MQ) traits are crucial in the practical breeding of malting barley, but the measurement of MQ traits is expensive and labor-intensive [14, 15]. In the whole process of brewing, numerous metabolic processes are involved, which results in distinct and time-dependent alterations in metabolite profiles [16]. Therefore, two previous study were conducted: The first study investigated the genetic variance in metabolomic profiles extracted from spring barley wort, where metabolomic features were found to be heritable and have great potential to be used in breeding for high MQ [17]; The second study investigated prediction of phenotypes using metabolomic features, where results showed that these phenotypic predictions were highly correlated to the actual phenotypes in a validation data set [18]. Keeping these two results in mind, the joint model for phenotypes and omics data proposed by Christensen et al. [12] may be the appropriate tool in practical barley breeding for the integration of both genomics and metabolomics into genetic evaluation.

Therefore, the aim of this study was to investigate the possibility of combining phenotypic, genomic, and metabolomic data for genetic evaluation of MQ traits in spring barley. To reach this goal, the joint model including both genomic and metabolomic information (MGBLUP) was compared with the baseline genomic model (GBLUP) to estimate variance components (VCs), and predict breeding values. Accuracies of predicted breeding values were evaluated from two cross-validation strategies: leave-one-year-out and leave-one-line-out schemes.

## Results

In this study, the VCs and heritabilities of MQ traits were first evaluated. Then the breeding values of MQ traits were predicted by using MGBLUP and GBLUP models, and assessed using two different cross-validation schemes.

### Descriptive statistics for malting quality traits

#### Error! Reference source not found

show descriptive statistics of all the MQ traits analyzed in this study. There were 2,430 records analyzed for five MQ traits: filtering speed (FS), extract yield (EY), wort color (WC), beta-glucan content (BG), and wort viscosity (WV). The averages of all the traits were 4.84 for FS, 82.61 for EY, 5.87 for WC, 217.98 for BG, and 1.47 for WV. The coefficient of phenotypic variance ranged from 2.23% for EY to 52.91% for BG.

**Table 1.**
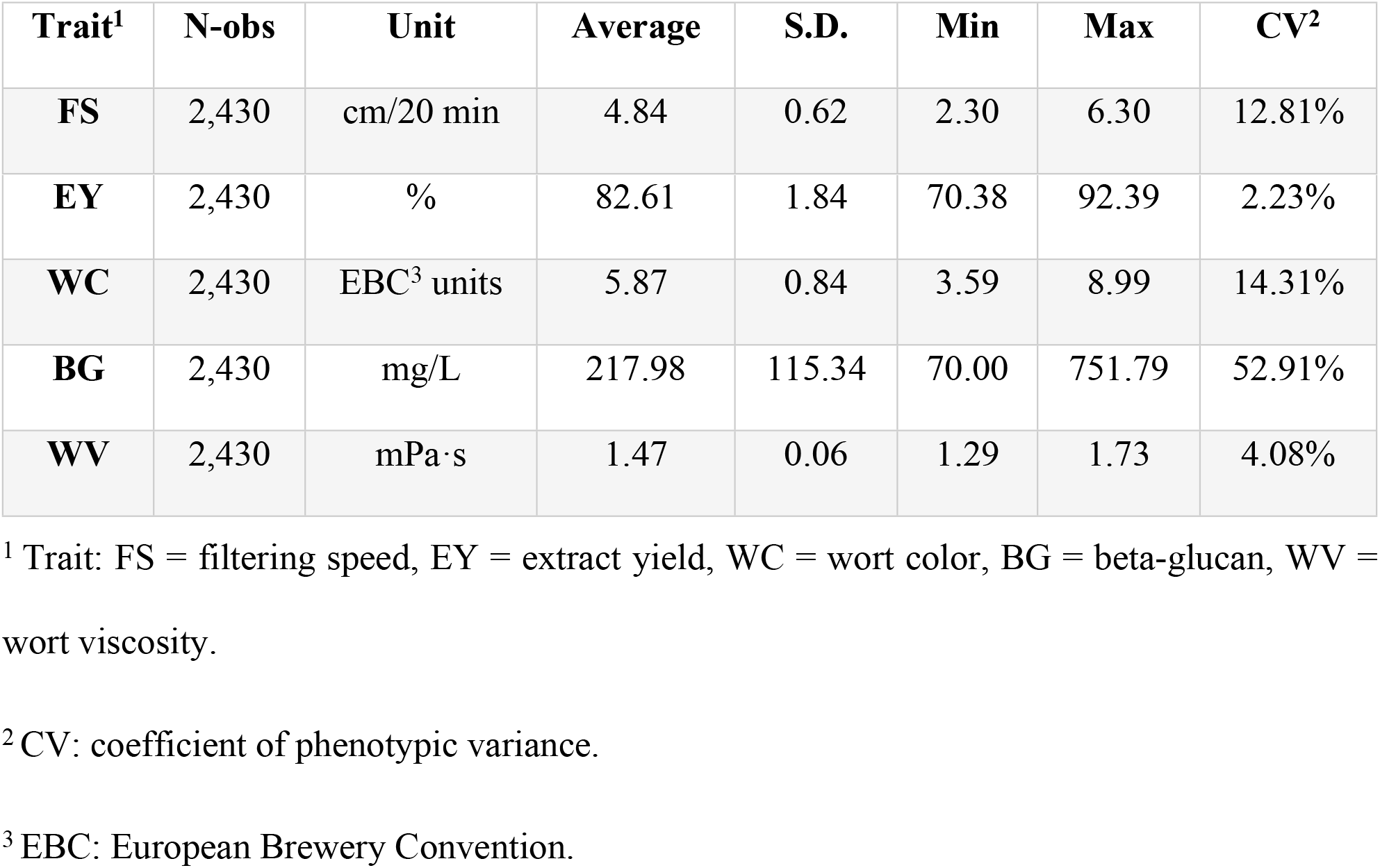
Descriptive statistics for malting quality traits.

| Trait<sup>1</sup> | N-obs | Unit | Average | S.D. | Min | Max | CV<sup>2</sup> |
| --- | --- | --- | --- | --- | --- | --- | --- |
| FS | 2,430 | cm/20 min | 4.84 | 0.62 | 2.30 | 6.30 | 12.81% |
| **EY** | 2,430 | % | 82.61 | 1.84 | 70.38 | 92.39 | 2.23% |
| **WC** | 2,430 | EBC<sup>3</sup> units | 5.87 | 0.84 | 3.59 | 8.99 | 14.31% |
| **BG** | 2,430 | mg/L | 217.98 | 115.34 | 70.00 | 751.79 | 52.91% |
| **WV** | 2,430 | mPa·s | 1.47 | 0.06 | 1.29 | 1.73 | 4.08% |

<sup>1</sup> Trait: FS = filtering speed, EY = extract yield, WC = wort color, BG = beta-glucan, WV = wort viscosity.
<sup>2</sup> CV: coefficient of phenotypic variance.
<sup>3</sup> EBC: European Brewery Convention.

### Estimated variance components

Estimates of variance components from GBLUP and MGBLUP are made comparable by dividing by the total phenotyping variance for each of the models. Relative variance component (RVC) is the percentage VC accounts for compared to the total phenotypic variance 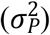 for each plot, and estimates of RVCs are shown in **Error! Reference source not found.** and supplementary S1 Table. The estimated genomic heritabilities 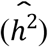, which are also RVCs for genomic effects (***g***), ranged from 0.060 (FS) to 0.279 (WC) when using GBLUP. When including metabolomics by using MGBLUP model, the 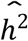 increased for all five MQ traits, and ranged from 0.123 (FS) to 0.283 (WC).

The 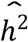 estimated from MGBLUP is the sum of an omics-mediated part and a residual part, and the decomposition of 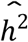 based on MGBLUP is listed in **Error! Reference source not found.**. As shown in **Error! Reference source not found.**, the residual 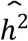 ranged from 0.003 (BG) to 0.069 (EY) and the omics-mediated 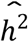 ranged from 0.113 (FS) to 0.270 (WC), showing that the omics-mediated 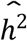 accounted for most of the 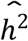.

Compared with GBLUP, the estimates of RVC for line effects (***ι***), and additive genotype by environment interaction effects (***i_g_***), from MGBLUP increased for FS and EY, but decreased for WC, BG and WV, while for residual effects (***e***), the trend was the opposite; RVC for line by environmental interaction effect (*i_ι_*) increased for FS, WC, WV, but decreased for EY and BG. Similarly, the trend for effects for batches of samples malted and mashed simultaneously (***t***) was opposite the trend for ***i_ι_***.

**Table 2.**
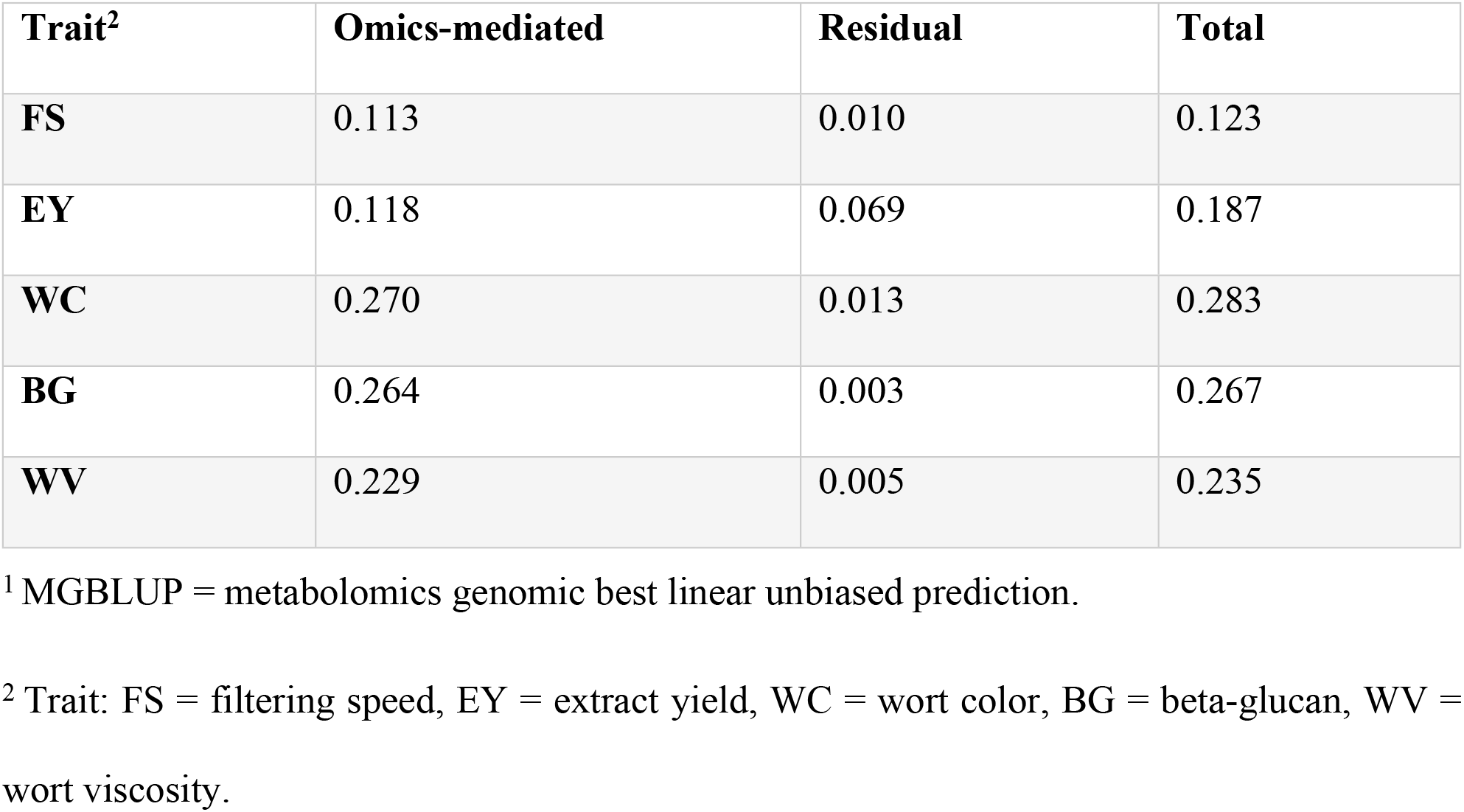
Heritability of malting quality traits using MGBLUP^1^.

| Trait<sup>2</sup> | Omics-mediated | Residual | Total |
| --- | --- | --- | --- |
| FS | 0.113 | 0.010 | 0.123 |
| EY | 0.118 | 0.069 | 0.187 |
| WC | 0.270 | 0.013 | 0.283 |
| BG | 0.264 | 0.003 | 0.267 |
| WV | 0.229 | 0.005 | 0.235 |

<sup>1</sup> MGBLUP = metabolomics genomic best linear unbiased prediction.
<sup>2</sup> Trait: FS = filtering speed, EY = extract yield, WC = wort color, BG = beta-glucan, WV = wort viscosity.

**Fig 1.**
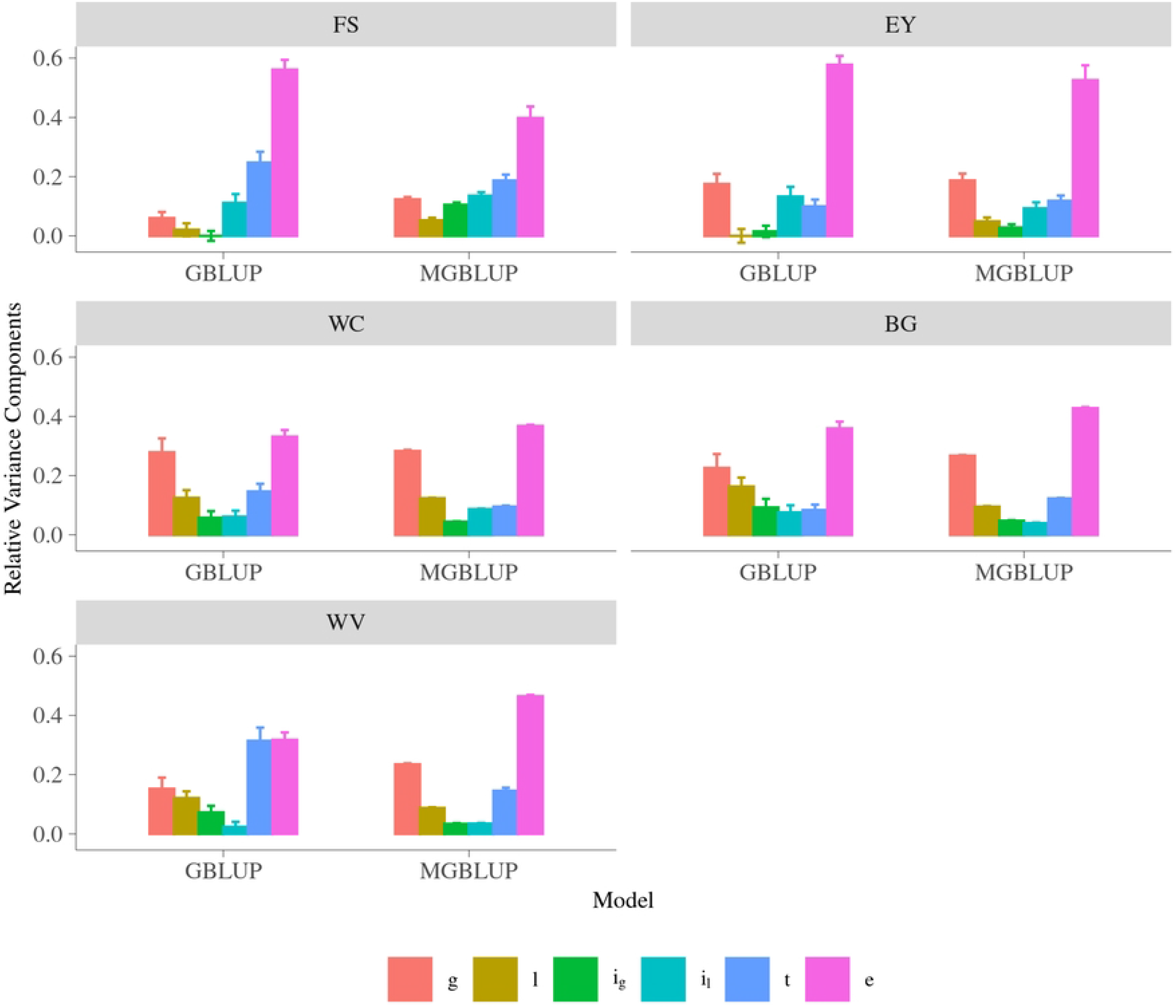
Proportion of total phenotypic variance explained by each component in malting quality traits. Trait: FS = filtering speed, EY = extract yield, WC = wort color, BG = beta-glucan, WV = wort viscosity; y-axis is relative variance component; g is relative variance of genomic effects, l is relative variance of line effect, i_g_ is relative variance of genotype by environmental effects, i_l_ is relative variance of line by environmental effects, t is relative variance of malt-mash effects, and e is relative variance of residuals. Method: GBLUP = genomic best linear unbiased prediction, MGBLUP = metabolomics genomic best linear unbiased prediction.

### Predicted breeding values

For GBLUP, predicted breeding values were computed by solving the GBLUP mixed model equations (MME) [19]. For MGBLUP, predicted breeding values were obtained by solving two MMEs successively, according to Christensen et al. [12]. In the first MME, effects of metabolomics and residual breeding value, i.e. genetic effects not mediated by metabolomics, on phenotype, were predicted. In the second MME, breeding values mediated by metabolomics, i.e. genetic effects on predicted metabolomic effects from the first MME, were predicted. The predicted total breeding value was the sum of these two predicted breeding values (mediated and not-mediated by metabolomics).

Accuracy of predicted breeding value is by definition the correlation between the predicted and the true breeding value, and this was assessed by dividing the whole dataset into a training data set and a validation data set, where predicted breeding values were computed based on the training data set, and validated against corrected phenotypes in the validation data set. In this study, prediction accuracy was the correlation between phenotypes corrected for fixed effects based on GBLUP and the predicted breeding values from each model, divided by square root of heritability estimated from each model in order to obtain an estimate of the correlation between predicted and true breeding values.

Two cross-validation schemes were applied in this study, and the accuracies of predicted breeding values are shown in **Error! Reference source not found.** for leave-one-year-out (LOYO) scheme and in **Error! Reference source not found.** for leave-one-line-out (LOLO) scheme. Accuracies obtained from MGBLUP model were higher than those from GBLUP model for all five traits in both cross-validation schemes (except for EY in LOLO which yielded same accuracy from both models).

**Fig 2.**
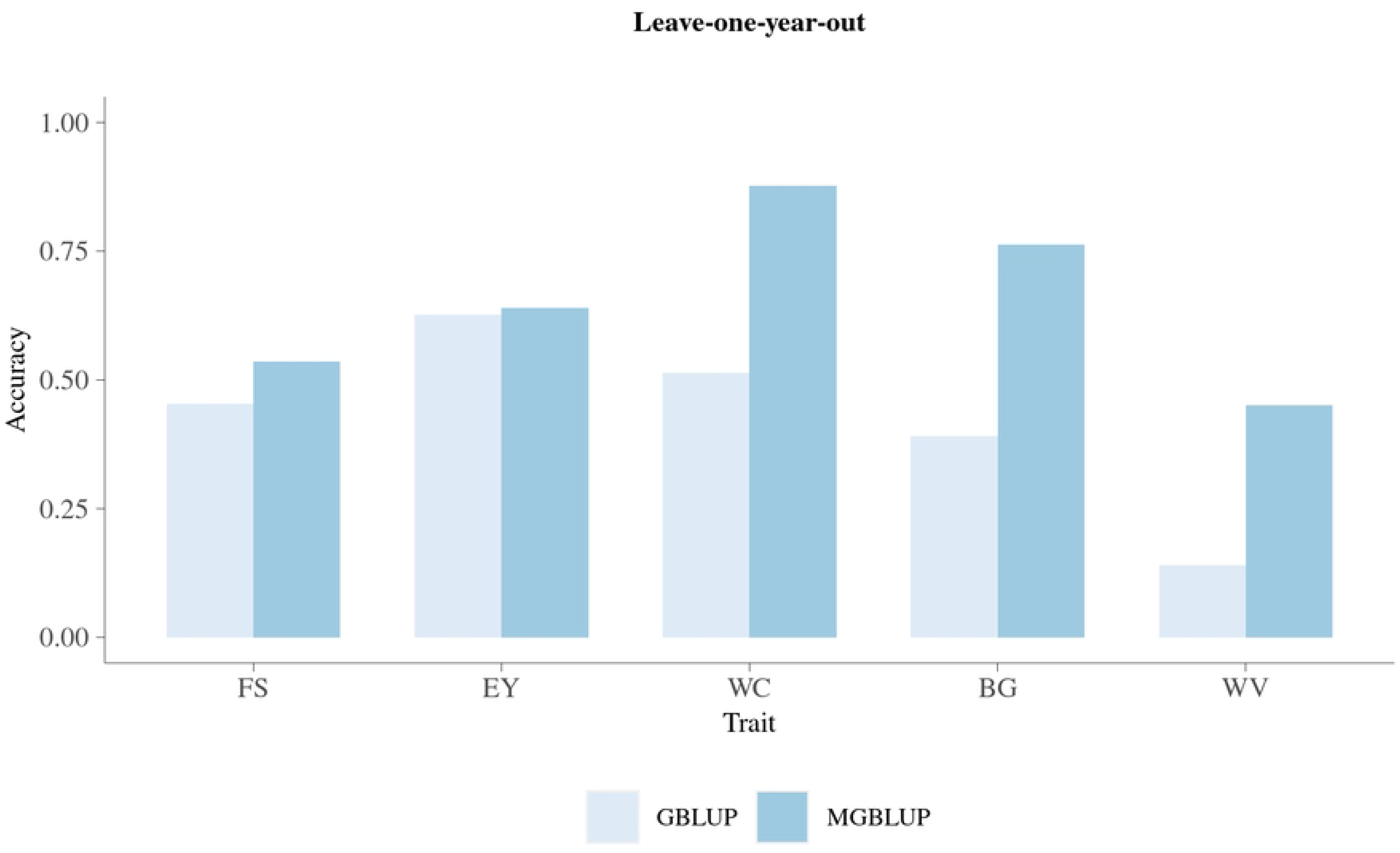
Accuracy of prediction for malting quality traits using leave-one-year-out cross-validation. Trait: FS = filtering speed, EY = extract yield, WC = wort color, BG = beta-glucan, WV = wort viscosity; y-axis is prediction accuracy. Method: GBLUP = genomic best linear unbiased prediction, MGBLUP = metabolomics genomic best linear unbiased prediction.

**Fig 3.**
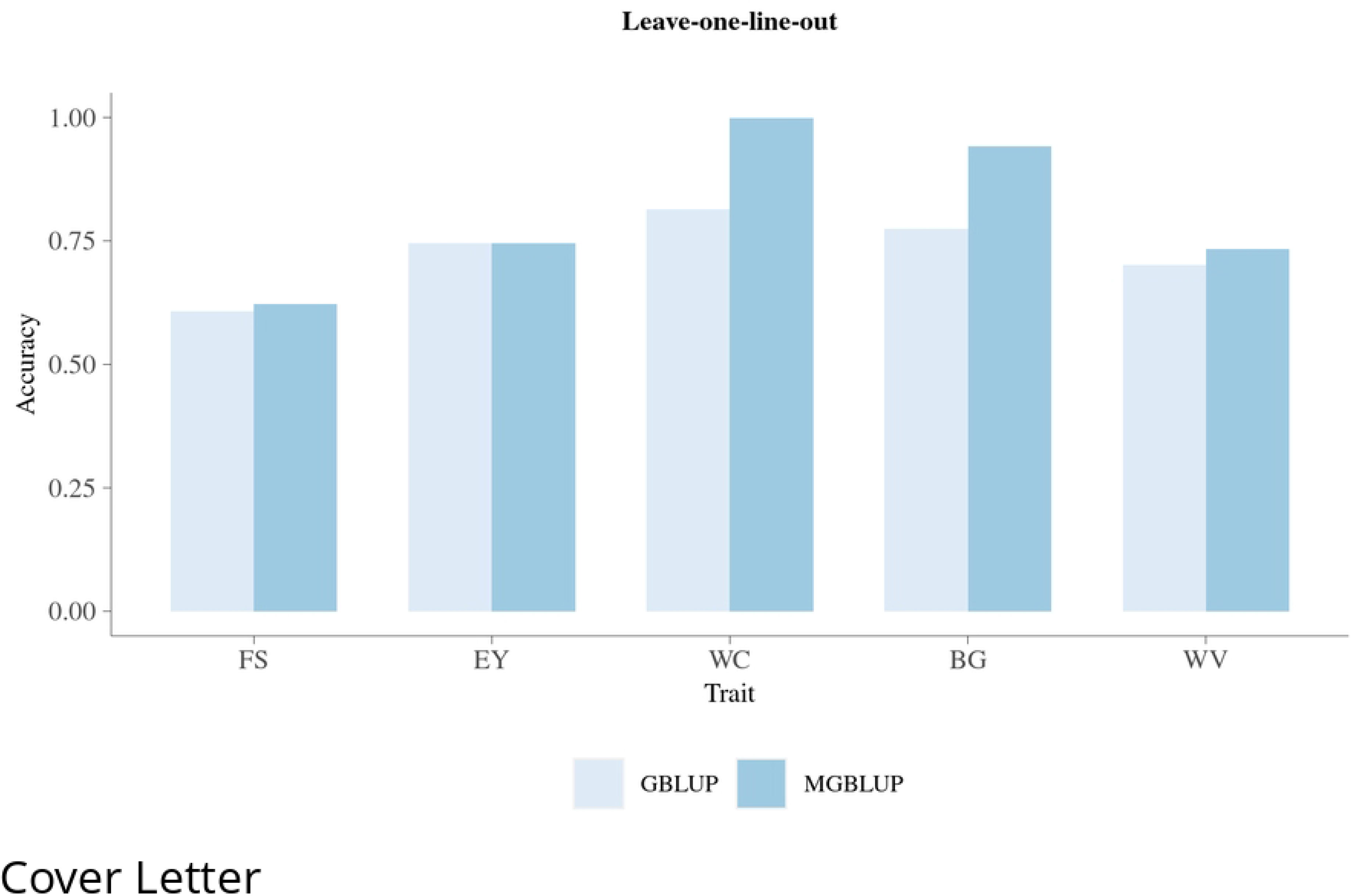
Accuracy of prediction for malting quality traits using leave-one-line-out cross-validation. Trait: FS = filtering speed, EY = extract yield, WC = wort color, BG = beta-glucan, WV = wort viscosity; y-axis is prediction accuracy. Method: GBLUP = genomic best linear unbiased prediction, MGBLUP = metabolomics genomic best linear unbiased prediction.

With LOYO scheme, the accuracy of prediction for breeding values ranged from 0.141 (WV) to 0.627 (EY) when using GBLUP, and increased to the range of 0.450 (WV) to 0.876 (WC) when using MGBLUP.

With LOLO scheme, the accuracy of predicted breeding values ranged from 0.607 (FS) to 0.814 (WC) when using GBLUP, and increased to the range of 0.623 (FS) to 0.998 (WC) when using MGBLUP.

## Discussion

Prediction of breeding values using 3,889 single-nucleotide polymorphism (SNP) markers and integrating metabolomic information with 24,018 metabolomic features (MFs) was carried out for a total of 2,430 plots of 562 spring barley malting lines, each phenotyped for five malting quality (MQ) traits. GBLUP and MGBLUP were compared. Accuracy of cross-validation was investigated by using leave-one-year-out (LOYO) and leave-one-line-out (LOLO) schemes.

Estimates of parameters were presented in the form of variance components relative to the total phenotypic variance (RVC), which made estimates from GBLUP and MGBLUP comparable. Increase in heritability was observed for all five traits of interests when including metabolomic information into the model. In MGBLUP, in the first step, residual heritability not captured by metabolomic effects was estimated in MGBLUP_1_ when fitting both genomic and metabolomic effects together. Then an omics-mediated heritability was estimated from MGBLUP_2_ when fitting genomic effects in the model with predicted metabolomics effects on phenotype as response variable. This way, two heritabilities are obtained from MGBLUP, an omics-mediated heritability, i.e. from genomic effects mediated by metabolomics, and a residual heritability, i.e. from genomic effects, not mediated by metabolomics, and the sum of the two heritabilies is the total heritability [12]. For all five traits investigated, the residual heritability accounted for only a small proportion of the total heritability, and the omics-mediated heritability accounted for a large proportion of the total heritability, which indicate that MFs play an important role in the biological path from gene to phenotype for these traits. This large proportion of omics-mediated heritability is also consistent with one of our previous studies, where significant genetic correlation was found between MFs and MQ traits [17]. The residual heritability of all five traits is smaller than the corresponding heritability estimated from GBLUP, which indicate that a part of the GBLUP heritability was distributed to the metabolomic effects in MGBLUP_1_, and then contributed to the omics-mediated heritability. The total heritability is larger for MGBLUP than heritability for GBLUP for all five traits, which may suggest that MGBLUP with two components of breeding value is better able to capture the genomic effects than GBLUP.

Two cross-validation schemes were carried out to evaluate two different aspects of prediction accuracy for MQ traits. In LOLO scheme, the aim was to predict breeding values of a line using phenotypes of other lines from the same breeding cycle, and hence the relationship between the training and the validation populations were stronger than in the LOYO scheme, and also the size of training population was larger than in the LOYO scheme. Therefore, it was expected that higher prediction accuracy could be obtained from LOLO than from LOYO, and the results obtained were consistent with this expectation. In addition, lines are generally much more related within year than across years. Therefore, in LOLO each line in the validation population has several close relatives in the training population, resulting in GBLUP having a high accuracy in this case, which means that those accuracies are difficult to improve further. The accuracies from LOLO have less practical importance than those from LOYO, because we usually want to predict one year ahead in the practical breeding.

The LOYO scheme is very relevant for a practical breeding system, since a new set of lines should be developed every year, and there is a large benefit of predicting breeding values of these lines based on metabolomics samples before the phenotypic records in replicated tests can be obtained. Therefore, LOYO scheme could be used to check if the model developed would be able to predict one breeding cycle from the other breeding cycles, as well as the ability that future crosses could be predicted based on current and previous crosses, which is very meaningful for practical barley breeding.

In both cross-validation schemes, the MGBLUP, integrating metabolomics and genomics in prediction, yielded higher accuracy of predicted breeding values than GBLUP. This means we can get clear improvements in accuracy when integrating metabolomics into genetic evaluation by using the model applied in this paper.

When using the LOYO scheme, a practical breeding program was mimicked. According to the prediction accuracy provided by GBLUP and MGBLUP models in this scheme, the five traits could be divided into two groups. The first group, consisting of FS and EY, had moderate improvement when applying MGBLUP compared with GBLUP model. While the second group, consisting of WC, BG and WV, had great improvement when applying MGBLUP compared with GBLUP. This larger improvement in the second group is consistent with the findings in our previous study, where much higher proportion of MFs had significant genetic correlation with the traits in second group (64.53% to 86.77%) than in the first group (6.09% to 11.88%) [17]. These results provide a clear indication that MGBLUP could be of great potential to be applied in the practical breeding for the traits in the second group.

In addition, from our results on heritability and prediction accuracy for the five traits from both GBLUP and MGBLUP, as expected, prediction accuracy generally increased with trait heritability [20]. Obviously, larger increases are possible for low heritability traits that have low prediction accuracy of GBLUP, since the maximum is 1.0.

The MGBLUP applied in this study integrated metabolomics by fitting a metabolomic similarity matrix built from MFs. Before building this matrix, the MFs were centered and standardized to a mean of 0 and standard deviation as 1 in order to equalize the MFs regardless of their intensities [21]. We tested the robustness of the model with respect to year effects, by comparing centering of the MFs, either across the three years or within each year. Centering within year theoretically minimize the variation caused by the factor of year compared with the centering across years, however, these different centerings yielded only small differences in parameter estimates or prediction accuracy, and the results reported are with the centering across years. These small differences indicate that the MGBLUP in the current study was able to handle the variation between years by fixed effects in the model. As the effect of environmental factors tend to be large in plant breeding, this is promising for the general robustness of the MGBLUP model.

The integration of multi-omics data has received great interests for predicting trait phenotypes in recent years [9–11]. In our previous study, we found that the prediction accuracy of prediction for MQ traits phenotypes using MFs was high [18]. However, the core of animal and plant breeding programs is the prediction of breeding values, and our study is the first implementation of a joint model for prediction of breeding values by integrating omics data into genetic evaluation. The successful implementation of such a joint model is very relevant to practical plant breeding where traits can be improved by selecting lines with large genetic potential. The MGBLUP model applied in the current study would also be applicable for integrating other omics data such as transcriptomics, and therefore it also could be interesting to expand the data resources integrated in the genetic evaluation of malting barley.

In conclusion, genetic evaluation of MQ traits in spring barley by MGBLUP provided very accurate prediction of breeding values, and MGBLUP is a robust approach for combining phenotypic, genomic and metabolomic data for prediction breeding values for MQ traits. We believe that the results from the current study have significant implications for practical breeding of barley and potentially many other species where accuracy of predicted breeding values is relatively low.

## Materials and Methods

All data used are available in a public accessible repository [22].

### Data

In this study, a total of 2,430 plots from 562 spring barley malting lines were included. These lines and the measurements of MQ traits were part of the standard breeding program from Nordic Seed A/S. Samples from two locations in Denmark were used, and they were taken from each plot individually and the data covered three years from 2014 to 2016. In both locations, the fields were divided into trials, which included 52 - 106 smaller plots (8.25 m^2^). Each trial was designed as a randomized complete block comprising 20 - 45 lines with three replicates of each line [23]. Each trial included two control lines in three replications. As a consequence, testing was conducted in a number of trials within each year-location combination. The malt sample from each plot was milled and extracted in water in order to produce a wort as described in a previous study [15]. The wort was used to measure five malting quality (MQ) traits which included filtering speed (FS), extract yield (EY), wort color (WC), beta-glucan content (BG), and wort viscosity (WV). Detailed description of these MQ traits can be found in a previous study [15]. Genotypic data were based on the Illumina iSelect9K barley chip and a total of 3,889 single-nucleotide polymorphism (SNP) markers were used after editing according to minor allele frequency more than 5% and missing markers less than 20%. Metabolomic features (MFs) were Nuclear Magnetic Resonance (NMR) data expressed as 24,018 intensities obtained from one-dimensional (1D) ^1^H NMR spectra, the intensities were integrated over small chemical shift intervals, expressed in parts per million (ppm) in the frequency range from 0.70 ppm to 9.00 ppm. The preparation of NMR for analysis is described in detail in our previous study [17].

### Statistical models and methods

A widely used approach for genetic evaluation is best linear unbiased prediction (BLUP). BLUP was derived by C. R. Henderson in 1950 [24, 25], and is a method for prediction of random effects in a statistical linear mixed model. The method has been widely used in animal and plant breeding for ranking of best individuals/lines in breeding programs. In this study, genomic BLUP (GBLUP) and metabolomic-genomic BLUP (MGBLUP) methods were compared.

### GBLUP

GBLUP is applied in this study as a baseline model, which is as below:

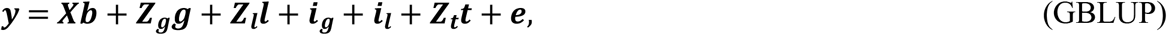

where ***y*** is the vector of records of each MQ trait, ***b*** is vector of location×year×trial effects to correct for differences caused by experimental location, year and trial, ***g*** is the vector of additive genomic effects for each line explained by genomic markers, ***ι*** is the vector of line effects for differences between lines not explained by additive effects of genomic markers, ***i_g_** is additive genotype by environment (six location×year environments) interaction effect which accounted for the additive genetic differences in genotype caused by various location×year environments, ***i_ι_*** is line by environmental interaction effect which accounted for the differences in line caused by various location×year environments but not explained by additive genotype by environment interaction effect, ***t*** is the vector of effects for batches of samples malted and mashed simultaneously which accounted for the environmental effects induced by the different batches, ***X, Z_g_, Z_ι_*** and ***Z_t_*** are the corresponding design matrices for ***b, g, l***, and ***t***, respectively, and ***e*** is vector of residual effects, i.e. variation that could not be explained by the other effects in the model. In this model, ***b*** is fixed effects parameter, and ***g, ι, i_g_, i_ι_, t*** and ***e*** are random effects with 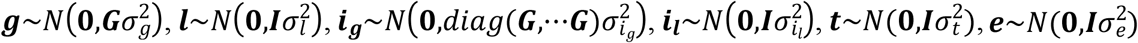, and these random effects are assumed to be independent of each other. Matrix ***G*** denotes the genomic additive relationship matrix computed using genomic data through VanRaden method 1 [26], matrix *di.ag*(***G***,...***G****) denotes the block-diagonal matrix with ***G*** as block-diagonal elements in six location×year environments, and ***I*** denotes the identity matrix.

### MGBLUP

MGBLUP is used in this study to allow the integration of both metabolomic and genomic data for prediction of breeding values. The model and methods closely follow the development by Christensen et al. [12]. First, we present the model in the context of our study, and second we explain the MGBLUP applied in this study.

The model developed by Christensen et al. [12] is a basic model to integrate different omics data into genetic evaluation. In our study, we extend this basic model to include additional random effects as in the GBLUP model, and also to incorporate the fact that we have multiple phenotypic and metabolomic records of each genotype. The model we use is a joint model for phenotypes and metabolomics intensities and specified by equations (1) and (2) below:

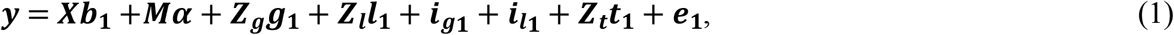

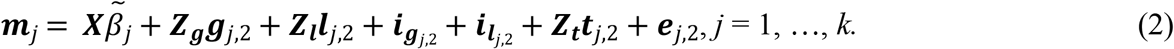

where ***y, X, b, Z_g_, Z_ι_, g, ι, i_g_, i_ι_, Z_t_, t, e*** are defined as for GBLUP model regardless of subscripts 1 or 2. Equation (1) describes the relationship between phenotypes and metabolomics intensities. Here, matrix ***M*** contains as columns the vectors ***m***_*1*_ to ***m**_k_* of metabolomics intensities for each of the *k* features, vector ***α*** contains regression effects of metabolomics intensities on phenotypes, and vector ***g***_1_ contains residual genetic effects, which are genetic effects on the phenotypes that are not mediated through the observed metabolomic intensities. Equation (2) for metabolomic feature *j* = 1, …, *k* describes the relationship between metabolomic intensities ***m***_*j*_ and genetic and environmental effects for the individuals. For the *j^th^* feature, vector *β_j_* contains fixed effects, ***g***_*j*,2_ is the vector of genetic effects on intensities, ***Λ***_*j*,2_ is the vector of line effects on intensities, ***i***_**g**_*j*,2__ is the vector of additive genotype by environment interaction effects on intensities, ***i***_***ι***_*j*,2__ is the vector of line by environmental interaction effects on intensities, ***t***_*j*,2_ is the vector of batch effects on intensities, and ***e***_*j*,2_ is vector of residual effects on intensities. All the random effect vectors are independent and their distributions are 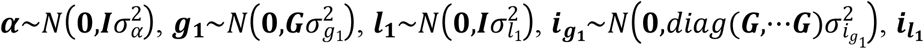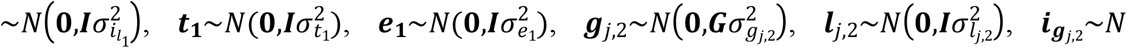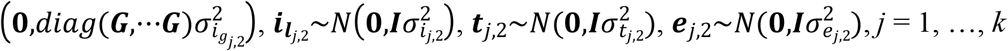.

In Christensen et al. [12], it was shown that predicted breeding values in the metabolomics-genomic model can be obtained by solving two mixed model equation systems successively, where each of these systems correspond to a linear mixed model. In our context this implies that inference in the Metabolomic-Genomic model can be obtained from successively applying MGBLUP_1_ and MGBLUP_2_ below.

The first step is:

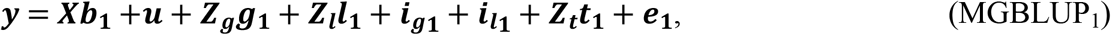

and the second step is:

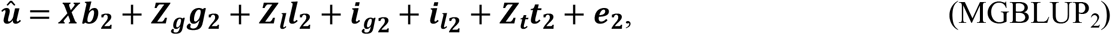

where ***u*** is the vector of metabolomic effects on phenotype and equal to ***Mα*** in equation (1), other effects in MGBLUP_1_ are the same as in equation (1), and effects in MGBLUP_2_ are defined similarly to those in equation (2). The metabolomics effects 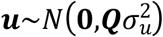, where 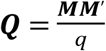 where ***M*** is a *p* × *q* matrix of adjusted, centered and scaled NMR intensities with *q* = 24,018 (equal to number of MFs) and *p* = 2,430 (equal to number of samples). In MGBLUP_2_, the 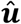 is the vector of predicted metabolomics effects from MGBLUP_1_, and the distribution of random effects are 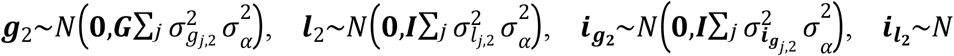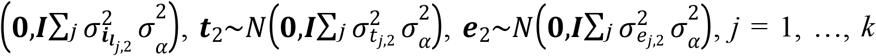. For MGBLUP, the predicted breeding values were calculated as the sum of predicted breeding values from MGBLUP_1_ and MGBLUP_2_ following the derivation in the pervious study [12],

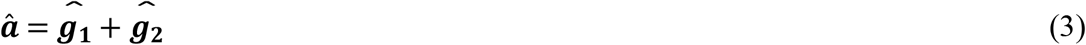

### Estimation of variance components

The full data set was used to estimate variance components (VCs) by the models described above. The VCs in all three models were estimated by restricted maximum likelihood (REML) using the DMU software package [27]. Relative variance component (RVC), which was the percentage each VC accounted for compared to the total phenotypic variance 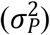 for each plot, was computed. The phenotypic variances were calculated as the sum of VCs in each model.

For GBLUP, 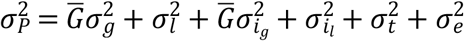, so that the genomic heritability of GBLUP is 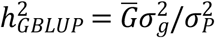, where 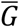 is the average of diagonal elements in the ***G*** matrix.

For MGBLUP, the calculation is 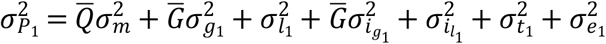, where 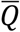 is the average diagonal of the ***Q*** matrix. So that the residual heritability is 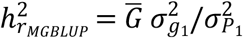 and the metabolomics variance ratio is 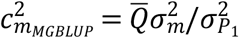. Furthermore, according to Christensen et al. [12], heritability of the metabolomics intensities can be obtained from MGBLUP_2_, i.e. here 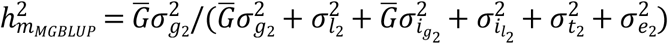, and therefore according to Christensen et al. [12], the genomic heritability based on the MGBLUP model is: 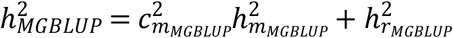.

It is common in plant breeding [28] to provide heritability of line mean instead of heritability of plot, as is done here. Such estimates for heritability of line mean will increase by introduction of better experimental design with more years and locations being tested, while heritability of plot reflects how heritable the traits of interests is in a more neutral way, and in addition, in our study the MFs were measured for each plot. Therefore, we chose to present the heritability of plot in this study.

### Cross-validation

Cross-validation was carried out to evaluate the accuracy of predicted breeding values. Two different cross-validation schemes, in which the whole dataset was divided into a training population (TP) and a validation population (VP), were investigated in this study based on two different hypotheses regarding factors that influence prediction accuracies. The two different schemes were leave-one-year-out (LOYO) and leave-one-line-out (LOLO). In LOYO, one out of three years was left out so that the accuracy of predicting one year from the other two years could be investigated. This scheme is similar to prediction of lines from a new breeding cycle. In LOLO, one out of 562 lines was left out so that the accuracy of predicting one line from all the other lines could be investigated. This scheme is similar to prediction of lines where no MQ phenotypes have been measured but other lines from the same breeding cycle have been MQ phenotyped.

In each round of the cross-validation, according to the setup of TP size, a certain number of the phenotypes were selected and the remaining phenotypes were masked. The phenotypes of the masked samples were predicted based on the TP together with the genomic and metabolomic information. Thereafter, the accuracy of prediction was calculated. Prediction accuracy was the correlation between phenotypes corrected for fixed effects based on GBLUP and the predicted breeding values from each model divided by square root of heritability estimated from each model. Calculated this way, the prediction accuracy is a measure of how well the predicted breeding values from the models predicted the phenotypes compared to the theoretical maximum given the proportion of variance that can be explained by additive genetic effects.

## Acknowledgments

The authors are grateful to Vahid Edriss, Nanna Hellum Kristensen, Jens Due Jensen, and Jette Andersen from Nordic Seed for field and laboratory work, Jihad Orabi and Hanne Svenstrup from Nordic Seed for SNP genotyping, Frans A. A. Mulder, Lars Alf Jensen and Benjamin Wahlqvist from Aarhus University for sample preparation and performing NMR measurements.

## Supporting information

**S1 Table. Relative variance components from two models for malting quality traits.**

Trait: FS = filtering speed, EY = extract yield, WC = wort color, BG = beta-glucan, WV = wort viscosity;

Method: GBLUP = genomic best linear unbiased prediction, MGBLUP = metabolomics genomic best linear unbiased prediction.

Effects: ***g*** is relative variance of genomic effects, ***l*** is relative variance of line effect, ***i_g_*** is relative variance of genotype by environmental effects, ***i_l_*** is relative variance of line by environmental effects, ***t*** is relative variance of malt-mash effects, and ***e*** is relative variance of residuals.

